# Characterisation of a highly diverged mitochondrial ATP synthase peripheral stalk subunit *b* in *Trypanosoma brucei*

**DOI:** 10.1101/2021.10.13.464200

**Authors:** Caroline E. Dewar, Silke Oeljeklaus, Bettina Warscheid, André Schneider

## Abstract

The mitochondrial F_1_F_o_ ATP synthase of *Trypanosoma brucei* has been studied in detail. Whereas its F_1_ moiety is relatively highly conserved in structure and composition, the same is not the case for the F_o_ part and the peripheral stalk. A core subunit of the latter, the normally conserved subunit *b*, could not be identified in trypanosomes suggesting that it might be absent. Here we have identified a 17 kDa mitochondrial protein of the inner membrane that is essential for normal growth, efficient oxidative phosphorylation and membrane potential maintenance. Pulldown experiments and native PAGE analysis indicate that the protein is associated with the F_1_F_o_ ATP synthase. Its ablation reduces the levels of F_o_ subunits, but not those of F_1_, and disturbs the cell cycle. HHpred analysis showed that the protein has structural similarities to subunit *b* of other species, indicating that the F_o_ part of the trypanosomal ATP synthase does contain a highly diverged subunit *b*. Thus, the F_o_ part of the trypanosomal ATPase synthase may be more widely conserved than initially thought.

## INTRODUCTION

Most cellular ATP is synthesised by the F_1_F_o_ ATP synthase complex which, in eukaryotic cells, is localised to mitochondria and plastids. The canonical mitochondrial F_1_F_o_ ATP synthase consists of two subcomplexes, the membrane-embedded F_o_ and the soluble F_1_ portions, which are connected by both a central stalk that rotates within the F_1_ moiety and a stationary peripheral stalk. The F_1_ moiety contains a hexamer of alternating α*-* and β*-*subunits surrounding a single γ-subunit, which functions as the central stalk. The γ subunit protrudes out from this headpiece to connect with the F_o_ moiety. The F_o_ moiety holds the proton translocation channel through the inner mitochondrial membrane (IM) formed by subunit *a* and a *c*-subunit oligomer in a ring conformation. The F_1_ subcomplex is generally conserved in terms of catalytic mechanism, inhibition of the reverse reaction by IF_1_, subunit stoichiometry, subunit sequences and structure. Proton movement through the F_o_ moiety rotates the *c*-ring and the central γ subunit, which induces asymmetrical conformational changes in the three active sites on the α*-* and β-subunit interfaces, resulting in the stepwise ATP synthesis cycle (1).

The catalytic hexamer is prevented from rotation with the central stalk and the *c* ring by the peripheral stalk, which acts as a stator. This is essential for the driving of conformational changes in the α_3_β_3_ hexamer and for the coupling of proton movement with ATP synthesis. In the yeast and mammalian enzymes, the conserved oligomycin sensitivity-conferring protein (OSCP), part of the peripheral stalk, sits at the pinnacle of the F_1_ head and interacts with the α-, *h*/F_6_ and *b* subunits (2–6). The central component of the peripheral stalk is subunit *b* which stretches from the OSCP to the membrane bound F_o_ moiety, and interacts with subunits *d* and *h*/F_6_ via stable coiled coils (5). There is also direct contact between a catalytic α subunit and subunits *b, d*, and *h*/F_6_ (7). In the membrane, subunit *b* interacts with subunits 8 and *d* (8, 9), the proton pore subunit *a* (6, 7, 9–11), subunits *f* (10), and the dimerisation subunits *e* and *g* (9).

Despite overall structural and functional conservation between F_1_F_o_ ATP synthases of all three domains (12), there is now evidence for substantial divergence between eukaryotic lineages in proteins making up the non-core parts of the complex (13–20). Subunits of the peripheral stalk and subunits involved in the anchorage of the peripheral stalk to the membrane, in holding the F_o_ *a* subunit in proximity to the *c*-ring and in the dimerisation domains all show in general low sequence conservation between species. However, while many non-core ATPase subunits show no sequence similarity between species, structural homology can sometimes still be detected (13, 14, 21).

Recently the unusual functioning of the mitochondrial F_1_F_o_ ATP synthase in the complex life cycle of the parasitic protozoan *Trypanosoma brucei* has attracted a lot of interest (14, 22). During the tsetse midgut stages of the life cycle, the complex generates ATP via oxidative phosphorylation using substrates derived from amino acid catabolism. Similar to other eukaryotes, trypanosomal F_1_F_o_ ATP synthase dimers are thought to facilitate cristae formation and may increase the efficiency of oxidative phosphorylation (23, 24). In the mammalian bloodstream forms of the parasite however, although still functioning in Fe/S cluster assembly, metabolism and ion homeostasis, the mitochondrion is reduced in volume and does not express a cytochrome-dependent electron transport chain or carry out oxidative phosphorylation (22). ATP is generated primarily through glycolysis, and unusually, the F_1_F_o_ ATP synthase acts in reverse as an ATPase, hydrolysing mitochondrial ATP to generate the essential membrane potential (ΔΨm) (25–29). Despite the importance of this role, the *T. brucei* mitochondrion is remarkably tolerant to both the level of F_1_F_o_ ATPase complexes present (30) and to alterations in the F_1_F_o_ ATPase structure (21, 25, 31).

The composition and structure of the F_1_F_o_ ATP synthase of procyclic *T. brucei* and of *Euglena gracilis* have been analysed in detail (13, 14, 23, 32–35). Trypanosomes and *Euglena* belong to the euglenozoans and thus are phylogenetically quite closely related. The architecture of the catalytic F_1_ moiety of both complexes from the two species is essentially canonical (34), containing the conserved subunits α, β, γ, δ and ε (33). Unusually, the α subunits of both are found cleaved into two chains (31, 36), an euglenozoan-specific feature of unknown biological relevance (37–39). The F_1_ headpiece also contains three copies of an additional euglenozoan-specific subunit, termed p18. This protein was first thought to be subunit *b* of the peripheral stalk (33, 39, 40), but has now been shown to be an F_1_ subunit that is associated with the α subunits. It facilitates the interaction between the F_1_ moiety and the peripheral stalk (34, 36).

In the F_o_ portion of these complexes, subunits ATPTB1, 3, 4, 6 and 12 are deemed to be euglenozoan-specific (13). There is to date no structural information on the F_o_ moiety including the peripheral stalk of the *T. brucei* enzyme complex. However, the structure of the peripheral stalk of F_1_F_o_ ATP synthase of *E. gracilis* has been defined and is highly unusual. It features an extended OSCP and a divergent *d* subunit homolog termed ATPTB2 that contacts the ATPTB3, ATPTB4 and p18 subunits to prevent futile F_1_ head rotation, substituting for a reduced subunit *b* (13).

BLAST analysis using the sequences of the peripheral stalk subunit *b* from either yeast, mammals or from *E. gracilis* as templates failed to identify a trypanosomal homolog, and it has therefore been speculated that the trypanosomal orthologue might be either highly reduced in size as in *E. gracilis* and therefore difficult to find, or that it might even be absent (14). Here we show by using a combination of biochemical, molecular, genetic and *in silico* analyses that the *T. brucei* F_1_F_o_ ATP synthase does contain an unusual, highly diverged subunit *b* that so far has been elusive.

## RESULTS

### Tb927.8.3070 is an integral mitochondrial inner membrane protein

Recently, we analysed the mitochondrial proteome of the parasitic protozoan *T. brucei* using the ImportOmics approach. The resulting proteome consisted of 1120 proteins, many of which are of unknown function and do not have orthologues outside the kinetoplastids (41). In the present study, we are focusing on one of these proteins, Tb927.8.3070, which is 145 aa in length and has a molecular mass of 17 kDa. Since the protein contains a single predicted transmembrane domain (TMD), but is not part of the previously characterised mitochondrial outer membrane (OM) proteome (42), it was assumed to be an integral inner membrane (IM) protein.

In order to test this prediction, we produced a transgenic *T. brucei* cell line allowing tetracycline-inducible expression of C-terminally myc-tagged Tb927.8.3070. Immunofluorescence (IF) analysis of tetracycline-induced cells using anti-myc antibodies indicated that myc-tagged Tb927.8.3070 co-localised with the mitochondrial marker atypical translocase of the outer membrane 40 (ATOM40) (Fig. 1A). In addition, cell extractions with 0.015% digitonin showed that myc-tagged Tb927.8.3070 co-fractionates with the voltage dependent anion channel (VDAC), which serves as another mitochondrial marker (Fig. 1B, left panel). The tagged protein was also exclusively recovered in the pellet when the crude mitochondrial fraction was subjected to carbonate extraction at high pH (Fig. 1B, right panel), along with the integral OM protein VDAC. This suggests that Tb927.8.3070 is an integral mitochondrial membrane protein consistent with its expected localisation in the mitochondrial IM, even though no N-terminal mitochondrial targeting sequence can be reliably predicted using multiple algorithms.

**Figure 1.**
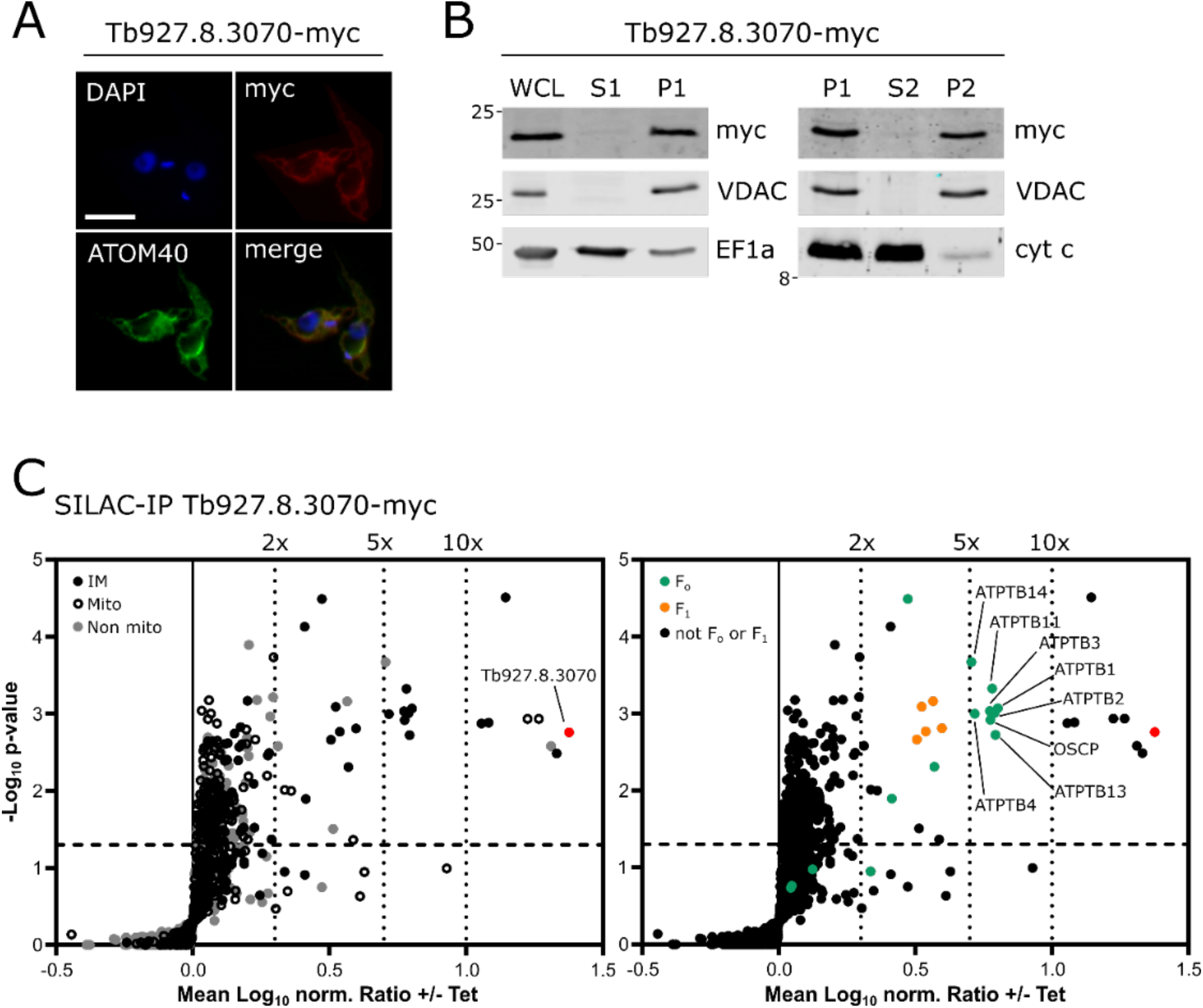
Tb927.8.3070 is an integral mitochondrial inner membrane protein that interacts with the F_1_F_o_ ATP synthase complex. **(A)** IFA image of a procyclic *T. brucei* cell line allowing tetracycline-inducible expression of Tb927.8.3070-myc. ATOM40 serves as a mitochondrial marker. DAPI marks both nuclear and mitochondrial DNA. Scale bar 10 μm. Cells were induced overnight. **(B)** Subcellular fractionation of cells expressing Tb927.8.3070-myc. Left panel: immunoblot analysis of whole cell lysates (WCL), digitonin-extracted mitochondria-enriched (P1) and soluble cytosolic (S1) fractions of cells expressing Tb927.8.3070-myc. The immunoblots were probed with anti-myc antibodies and antisera against VDAC and elongation factor 1-alpha (EF1a), which serve as markers for mitochondria and cytosol, respectively. Right panel: digitonin-extracted mitochondria-enriched fractions (P1) were subjected to alkaline carbonate extraction performed at pH 11.5 resulting in membrane-enriched pellet (P2) and soluble supernatant (S2) fractions. Subsequent immunoblots were probed with anti-myc and antisera against VDAC and cytochrome *c* (cyt *c*), which serve as markers for integral and peripheral membrane proteins, respectively. **(C)** Volcano plots of a SILAC-IP analysis of crude mitochondrial extracts from Tb927.8.3070-myc expressing cells. Cells were induced with tetracycline for 1 day. Proteins were quantified in three independent biological replicates, with the mean log_2_ of ratios (+/-Tet) plotted against the −log_2_ *P* value (two-sided *t*-test). The bait is shown in red. The horizontal dashed line shows a significance level of p = 0.05. The vertical dotted lines mark specified enrichment factors. Left panel, the following groups of proteins are highlighted: mitochondrial IM proteins (IM, black), other proteins of the mitochondrial importome (Mito, black with a white centre) and non-mitochondrial proteins (Non mito, grey). Right panel, the following groups of proteins are highlighted: ATP synthase F_o_ subunits (F_o_, orange) and F_1_ subunits (F_1,_ green).

### Tb927.8.3070 interacts with the F_1_F_o_ ATP synthase complex

To analyse whether Tb927.8.3070 was contained within a protein complex, a transgenic *T. brucei* cell line allowing inducible expression of Tb927.8.3070-myc was subjected to stable isotope labelling by amino acids in cell culture (SILAC) followed by an anti-myc immunoprecipitation (IP) from mitochondria-enriched fractions. The resulting eluates were analysed by quantitative mass spectrometry (MS). Including the myc-tagged Tb927.8.3070, which served as bait, 30 proteins were found significantly more than 2-fold enriched in this IP (Fig. 1C, Table S1). 25 of these proteins were members of the mitochondrial importome (41) (Fig. 1C, left panel), with 19 designated as being mitochondrial IM proteins (42). Strikingly, 16 previously identified subunits of the F_1_F_o_ ATP synthase (33) were significantly enriched more than 2-fold (Fig. 1C, right panel), suggesting that tagged Tb927.8.3070 might be a component of or interact with the F_1_F_o_ ATP synthase. Interestingly, the F_1_F_o_ ATP synthase subunits that were found most highly enriched (between 5 to 10-fold) included OSCP and ATPTB2, known subunits of the peripheral stalk, and ATBTB1, 3, 4, 11, 13 and 14 of the F_o_ moiety.

Other proteins which were found significantly more than 10-fold enriched include a M76 peptidase with homology to yeast and mammalian ATP23, which is a protease of the IM required for F_1_F_o_ ATP synthase assembly (43, 44), phosphatidylserine decarboxylase (PSD) (45), and cardiolipin-dependent protein 17 (CLDP17) (46). The remaining four proteins are kinetoplastid-specific proteins of unknown function (Fig. S1).

### Tb927.8.3070 is an essential protein and co-migrates with the F_1_F_o_ ATP synthase monomer

To investigate the function of Tb927.8.3070, we produced a tetracycline-inducible RNAi cell line of procyclic *T. brucei*. Ablation of Tb927.8.3070 caused a growth retardation by day 2 post induction (Fig. 2A). Next, we analysed the consequences of Tb927.8.3070 ablation on the F_1_F_o_ ATP synthase complex and subcomplexes. Blue native polyacrylamide gel electrophoresis (BN-PAGE) and subsequent immunoblot analysis using an antibody against F_1_ ATP synthase subunit p18 (33, 47, 48) in Tb927.8.3070-RNAi cells showed that the levels of F_1_F_o_ dimer and monomer were steadily decreased upon Tb927.8.3070 depletion to less than around 40% of the levels seen in uninduced cells by day 3 post induction (Fig. 2B). Interestingly, a similar, albeit stronger, phenotype was previously seen upon depletion of F_o_ subunit ATPTB1 and peripheral stalk subunits OSCP and ATPTB2 (21, 30, 33). In contrast, the levels of the free F_1_ moiety doublet were increased up to 4-fold by day 3 after Tb927.8.3070-RNAi induction (Fig 2B). This doublet likely represents the core F_1_ subcomplex with and without *c* ring attached, as in mammalian cells (21, 33, 49). These results are similar to those previously obtained after OSCP, ATPTB1 and ATPTB2 depletion (30, 33), and contrast with the effect of F_1_ subunit depletion, which reduces the level of both F_1_F_o_ and F_1_ ATP synthase complexes (33, 36).

**Figure 2.**
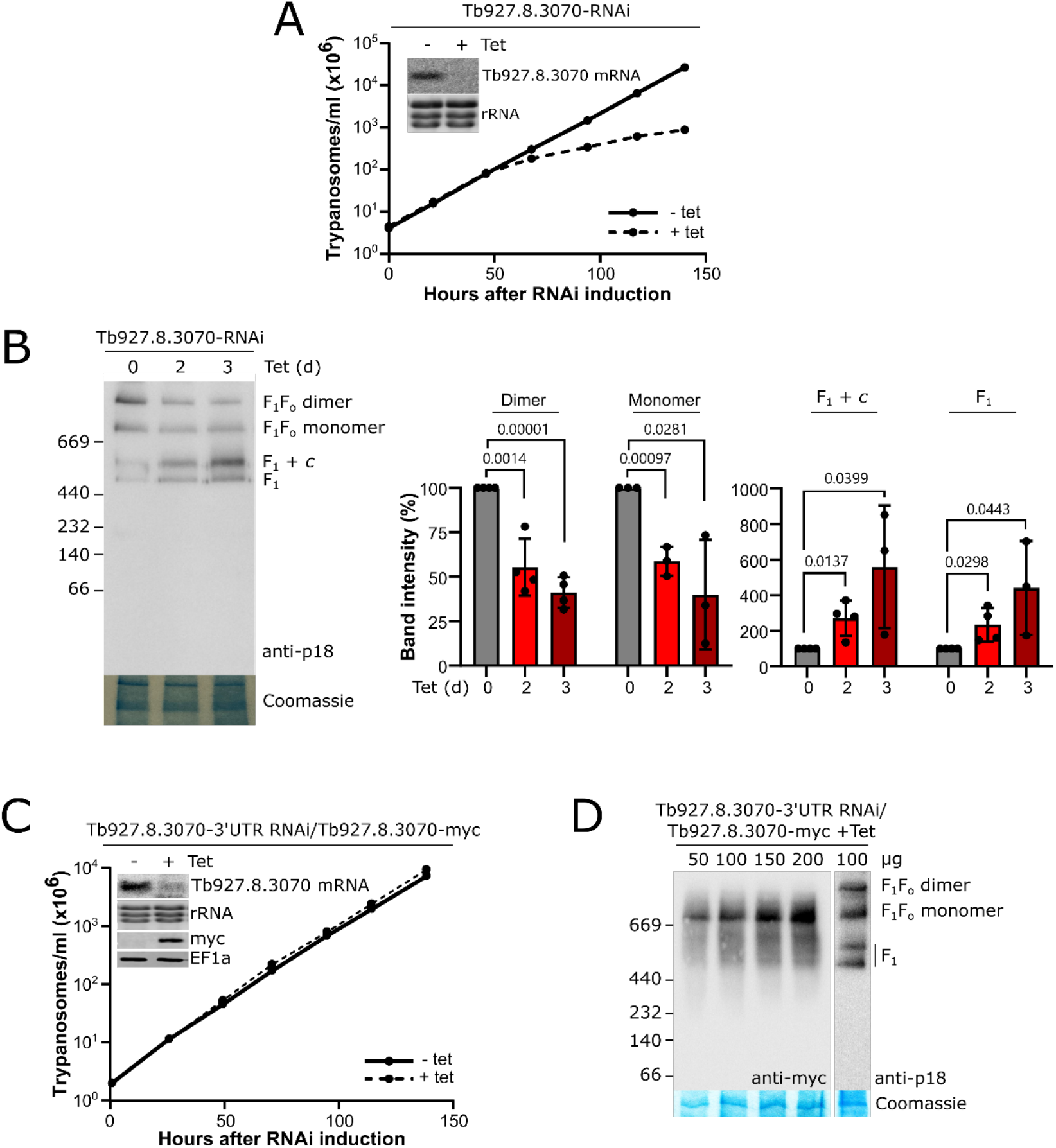
Tb927.8.3070 is an essential protein and co-migrates with the F_1_F_o_ ATP synthase monomer. **(A)** Growth curve analysis of a procyclic Tb927.8.3070-RNAi cell line. Analysis was performed in triplicate, with error bars showing standard deviation (too small to be visible). Inset, Northern analysis after 2 days of induction of the level of Tb927.8.3070 mRNA. EtBr-stained rRNAs acts as a loading control. **(B)** Left panel, BN PAGE analysis of crude mitochondrial extracts of Tb927.8.30-RNAi cells. ATP synthase complexes were visualised with a polyclonal antibody against ATP synthase F_1_ subunit p18. A section of the Coomassie stained gel serves as a loading control. Right panel, quantification of the BN PAGE using ImageJ. Error bars indicate standard deviation of three or four repeat experiments. P-values were calculated using an unpaired t test. **(C)** Growth curve analysis of the Tb927.8.3070-RNAi 3’ UTR cell line expressing Tb927.8.3070-myc. This cell line targets the 3’ UTR of Tb927.8.3070 and therefore allows reexpression of the protein in a different genomic context. Analysis was performed in triplicate, with error bars showing standard deviation. Inset, Northern analysis after two days of induction of the level of Tb927.8.3070 mRNA and the corresponding immunoblot analysis of the Tb927.8.3070-myc levels. EtBr stained rRNA species and EF1a act as a loading controls. **(D)** BN PAGE analysis of crude mitochondrial extracts from cells used in C induced for 2 days, with loading as indicated. The complexes indicated were visualised as in (B).

To exclude off-target effects in the Tb927.8.3070-RNAi cell line and to demonstrate that the tagged Tb927.8.3070 is functional, a cell line was generated where Tb927.8.3070 was downregulated by targeting its 3’UTR for RNAi upon tetracycline induction while a C-terminally myc-tagged version of Tb927.8.3070 was ectopically expressed. Fig. 2C shows that expression of Tb927.8.3070-myc fully restored growth of the induced RNAi cell line. BN-PAGE analysis of crude mitochondrial extracts from this induced cell line revealed the characteristic four band pattern of the *T. brucei* F_1_F_o_ ATP synthase that could be detected by an antibody specific for p18 (Fig. 2D, right lane). When the same samples were probed with anti-myc antibodies, only the F_1_F_o_ ATP synthase monomer was detected (Fig. 2D, left lanes), suggesting that the C-terminal myc-tag of Tb927.8.3070 interferes with F_1_F_o_ ATP synthase dimer formation.

In summary, these results show that depletion of Tb927.8.3070 has a similar effect to the depletion of ATPB1, OSCP and ATPB2 (21, 30, 33) and therefore is presumably not part of the F_1_ moiety but rather a subunit of the F_o_ ATP synthase subcomplex or the peripheral stalk associated with it.

### Tb927.8.3070 is a putative homolog of F_1_F_o_ ATP synthase subunit *b*

Interestingly, the core component of the peripheral stalk, subunit *b*, has not been identified in *T. brucei*. This is somewhat surprising since the composition of trypanosomal F_1_F_o_ ATP synthase has been analysed in detail using proteomics (33), and a highly diverged subunit *b* had been found in the phylogenetically related *E. gracilis* (13). However, reciprocal BLAST searches using either the *E. gracilis* subunit *b* or Tb927.8.3070 as templates did not retrieve either protein or any F_1_F_o_ ATP synthase subunit of other species. It should be taken into consideration though that the two proteins are very small, 17 kDa for Tb927.8.3070 and 12.7 kDa for *Euglena* subunit *b*, which would make it difficult to detect homology of diverged sequences. In fact, without structural information on the peripheral stalk it would not have been possible to identify the highly diverged subunit *b* of *E. gracilis*.

Thus, we wanted to investigate the possibility that Tb927.8.3070 might be the F_o_ ATP synthase subunit *b*. We used the HHpred algorithm that is based on the comparison of profile hidden Markov models (HMMs) which often is able to establish connections to remotely homologous, characterised proteins (50). The HHpred analysis retrieved 45 proteins of known function which shared low sequence and structural similarity with Tb927.8.3070 (Fig. S2A). 11 of these had regions of significant structural similarity of more than 80 amino acids. Most interestingly, four of them were related to mitochondrial ATP synthase subunit *b*: the spinach chloroplast subunit atpF, the yeast subunit ATP4, and the atpF subunits of two bacterial species, *Mycobacteria* and *Bacillus* (Fig. 3A, Fig. S2A highlighted in blue, S2B). Strikingly, in all four cases the region showing similarity with Tb927.8.3070 has the same relative position, including the experimentally confirmed TMDs and an approximately 60 aa C-terminal flanking sequence of the four F_1_F_o_ ATP subunit *b* proteins (Fig. 3A). Tb927.8.3070 is a shorter protein, with the spinach, yeast and bacterial proteins having a longer C-terminal extension past the region of structural similarity. BLAST detects Tb927.8.3070 sequence homologs only in kinetoplastids (Fig. S2C). Thus, based on these bioinformatic analyses and on our experimental results, we propose that Tb927.8.3070 is a highly divergent form of subunit *b* of the peripheral stalk of the trypanosomal F_1_F_o_ ATP synthase.

**Figure 3.**
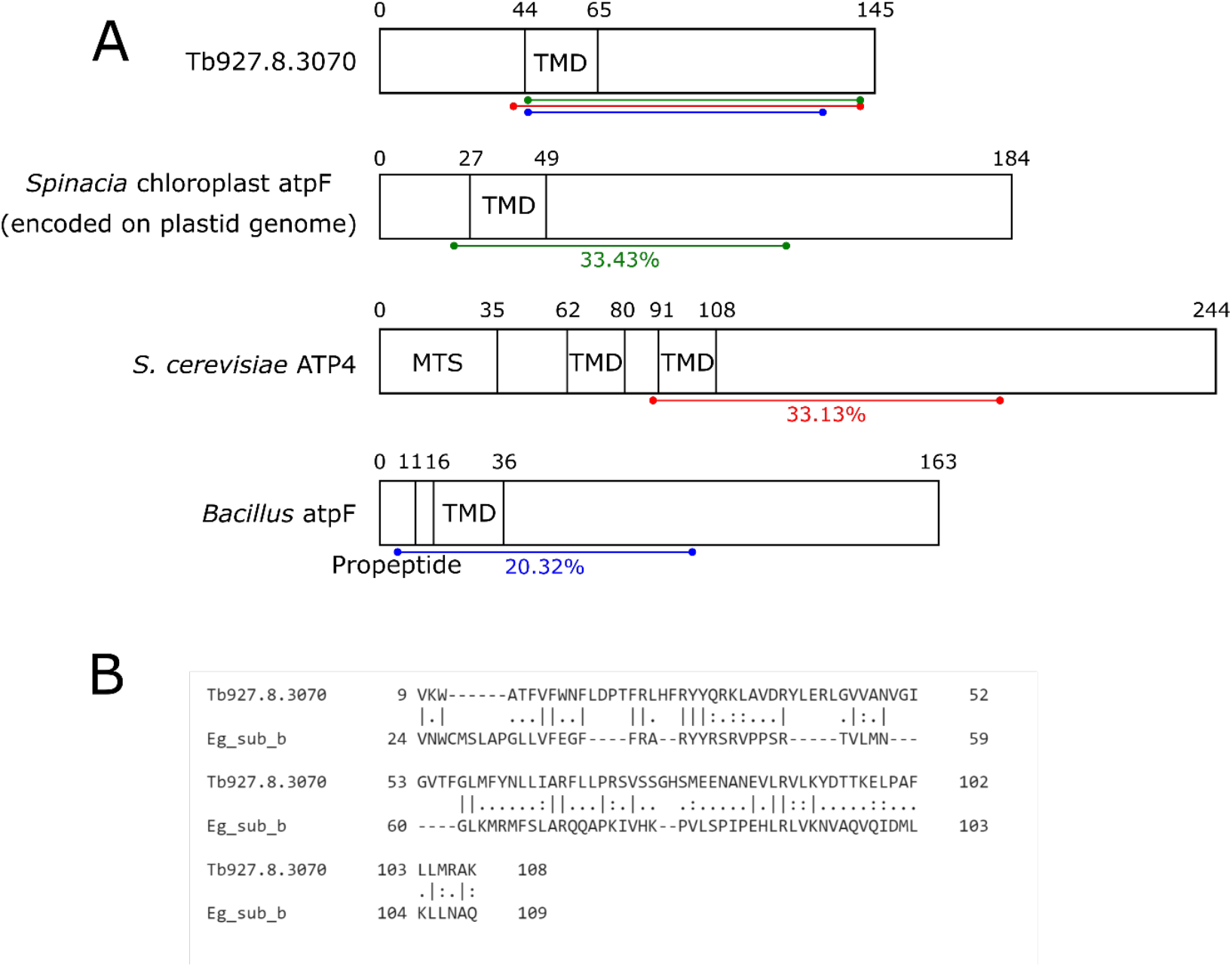
Tb927.8.3070 is a homolog of F_1_F_o_ ATP synthase subunit *b*. **(A)** A schematic showing a comparison of domains predicted in Tb927.8.3070 and the homologs of ATP synthase subunit *b* in spinach chloroplasts (atpF), yeast (ATP4) and Bacillus species (atpF). The regions of secondary structure homology as determined by HHpred are indicated by the coloured lines. Transmembrane domains (TMD) were predicted by HMMTOP and TMPred (74, 75). The yeast ATP4 mitochondrial targeting sequence (MTS) was previously defined experimentally (76). **(B)** An alignment of the protein sequences of Tb927.8.3070 and *E. gracilis* subunit *b* using the EMBOSS Water Pairwise Sequence Alignment tool (77).

It is somewhat surprising that HHpred analysis of the trypanosomal subunit *b* did not retrieve its counterpart in the relatively closely related *Euglena*. However, a pairwise alignment of the *E. gracilis* subunit *b* protein sequence with the trypanosomal protein showed that a region covering the central 70% of Tb927.8.3070 has 23.6% identity to the *E. gracilis* protein, suggesting that the proteins do share some limited sequence conservation (Fig. 3B). While this similarity is low, there are homologous F_o_F_1_ ATP synthase subunits which share less sequence identity between *Euglena* and *T. brucei*, indicating that the observed similarity might indeed be due to common descent (14).

### Subunit *b* ablation selectively depletes F_o_ ATP synthase subunits

Next, we used a SILAC-MS approach to determine the consequences of subunit *b* depletion on the mitochondrial proteome. After 3 days of RNAi induction, mitochondria-enriched fractions of uninduced (control) and induced SILAC Tb927.8.3070 RNAi cells were prepared and analysed by quantitative MS. We found 14 subunits of the F_o_ moiety of the F_1_F_o_ ATP synthase complex significantly decreased upon subunit *b* depletion (Fig. 4A, green), with 10 of these subunits (ATPTB1, 2, 5-7, 10-13) more than 1.5-fold depleted (Table S2). However, the levels of all six subunits of the F_1_ moiety (α, β, γ, δ, ε and p18) were unaffected in the same experiment (Fig. 4A, orange). This is similar to what was observed upon knockdown of the F_o_ subunit ATPB1 or the peripheral stalk subunits OSCP and ATPB2 (21, 30, 33), and in contrast to the knockdown of the F_1_ subunits α, β and p18, where the stability of both F_1_ and F_o_ components were affected (25, 26, 33). There were seven proteins not annotated as ATP synthase subunits found to be significantly more than 1.5-fold depleted in this experiment (Fig. 4B): the axonemal inner arm dynein light chain (51), three trypanosomatid-specific proteins of unknown function that are found in the mitochondrial importome (Tb927.6.590, Tb927.2.5930 and Tb927.10.9120) and three trypanosomatid-specific proteins of unknown function not detected in the importome (Tb927.9.7980, Tb927.10.1430 and Tb927.11.9940).

**Figure 4.**
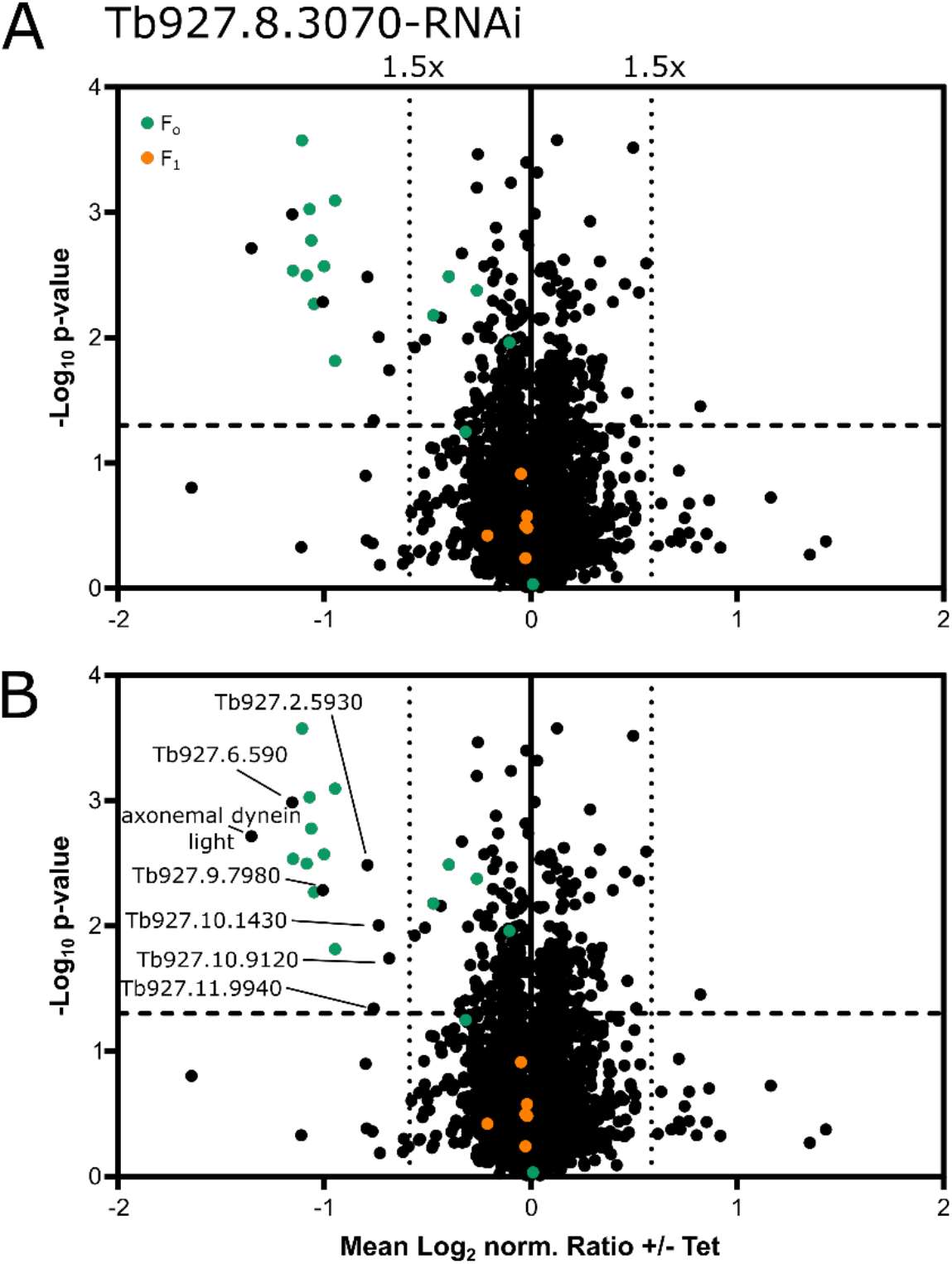
Tb927.8.3070 ablation selectively depletes F_o_ ATP synthase subunits. **(A)** Volcano plots of a SILAC-based quantitative MS analysis of crude mitochondrial extracts of uninduced and induced Tb927.8.3070-RNAi cells. Cells grown in SILAC media were harvested after 4 days of induction. Proteins were quantified in three independent biological replicates, with the mean log_2_ of normalised (norm.) ratios plotted against the −log_2_ *P* value (two-sided *t*-test). Tb927.8.3070 the RNAi target was not detected in this analysis. The horizontal dashed line indicates a significance level of p = 0.05. The vertical dotted lines mark proteins with a 1.5-fold change in abundance compared to control cells. The following groups of proteins are highlighted: ATP synthase F_o_ subunits (F_o_, orange) and F_1_ subunits (F_1,_ green). **(B)** As in (A), but proteins downregulated more than 1.5-fold are labelled with either their name or accession numbers.

In summary, these data are consistent with those shown in Fig. 2B and 2D, as well as with the HHPred analysis shown in Fig. 4 and indicate that Tb927.8.3070 is indeed the missing subunit *b*.

### Ablation of subunit *b* affects mitochondrial physiology

Mitochondrial ATP production by oxidative phosphorylation requires both intact F_o_ and F_1_ moieties of the ATP synthase. In order to determine the role of subunit *b* in this process, we performed *in organello* ATP production assays using digitonin-extracted crude mitochondrial fractions of the uninduced and induced RNAi cell line (52). It had previously been shown that in such assays depletion of the F_o_ subunit ATPTB1 or the peripheral stalk subunit ATPTB2 strongly impeded ATP production by oxidative phosphorylation using succinate as a substrate (33). When we measured the effect of subunit *b* depletion on ATP production levels, we found a significant decrease in the succinate-mediated ATP production of around 50% by day 3 post induction (Fig. 5A). This decrease in activity is specific to oxidative phosphorylation, as the level of ATP produced by substrate level phosphorylation stimulated by either α-ketoglutarate or pyruvate was unaffected. Subunit *b* thus is required for efficient oxidative phosphorylation in mitochondria of procyclic cells.

**Figure 5.**
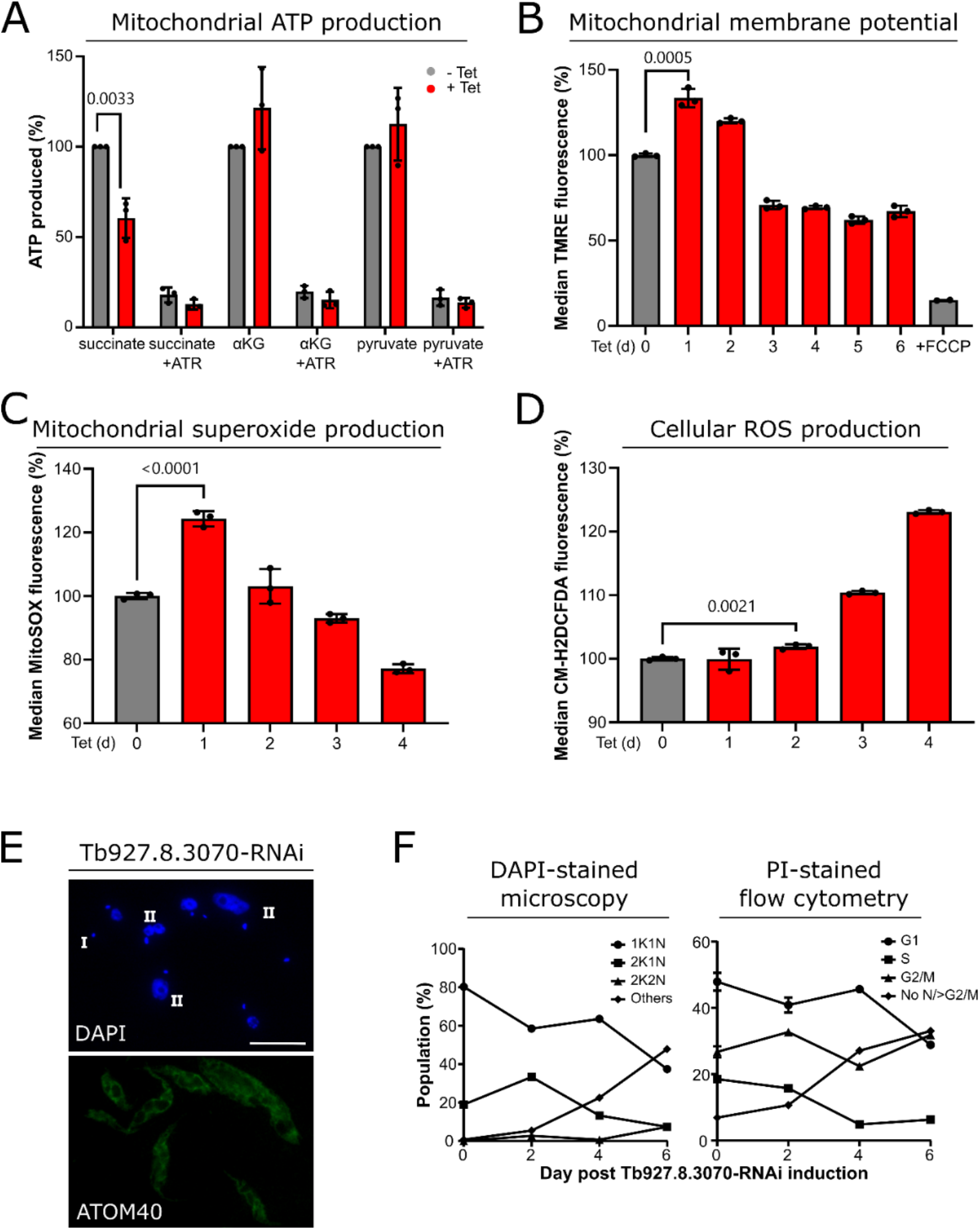
Tb927.8.3070 ablation affects mitochondrial physiology. **(A)** ATP production analysis of crude mitochondrial lysates from uninduced and induced Tb927.8.3070-RNAi cells. Oxidative phosphorylation or substrate level phosphorylation was induced by the addition of either succinate, α-ketoglutarate (aKG) or pyruvate plus succinate (pyruvate) as substrates. Atractyloside (ATR), an inhibitor of the ATP/ADP translocator selectively prevents mitochondrial ATP production. Three biological replicates were analysed, with error bars showing standard deviation. The p-value was calculated by an unpaired t test. The mean of ATP production detected in crude mitochondrial lysates of uninduced cells treated with the respective substrate was set to 100%. **(B)** Measurement of mitochondrial ΔΨm in the uninduced and induced Tb927.8.3070-RNAi cell line. Analysis was performed by measuring TMRE fluorescence using flow cytometry. Three biological replicates were analysed, with error bars showing standard deviation. The p-value was calculated using an unpaired t test. The average TMRE fluorescence of the uninduced cell line was set to 100%. Addition of FCCP, a mitochondrial membrane potential uncoupler, acts as a negative control. **(C)** Measurement of mitochondrial superoxide production in uninduced and induced Tb927.8.3070-RNAi cell lines. Analysis was performed by measuring MitoSOX fluorescence using flow cytometry. Three biological replicates were analysed, with error bars showing standard deviation. The p-value was calculated using an unpaired t test. The average MitoSOX fluorescence of the uninduced cell line was set to 100%. **(D)** Measurement of cellular reactive oxygen species production in the uninduced and induced Tb927.8.3070-RNAi cell line. Analysis was performed by measuring CM-H_2_DCFDA fluorescence using flow cytometry. Three biological replicates were analysed, with error bars showing standard deviation. The p-value was calculated using an unpaired t test. The average CM-H_2_DCFDA fluorescence of the uninduced cell line was set to 100%. **(E)** Representative IF image of Tb927.8.3070-RNAi cells induced for 4d. Top panel, DAPI marks both nuclear and mitochondrial DNA. Bottom panel, ATOM40 serves as a mitochondrial marker. Scale bar 10 μm. **(F)** Analysis of cell cycle progression upon depletion of Tb927.8.3070 by RNAi. Left panel, the visualisation of DAPI-stained cells (n>130 cells per time point). Right panel, the quantification of PI fluorescence using flow cytometry. The proportion of cells found in each cell cycle stage are shown as a percentage of the total population. Flow cytometry analysis was performed in three biological replicates. The average % value here are presented with error bars showing standard deviation.

As in other systems, the F_1_F_o_ ATP synthase of procyclic *T. brucei* utilises the ΔΨm generated by the mitochondrial electron transport chain to power ATP production. One known side effect of reduced ATP synthase activity in the PCF is the hyperpolarisation of the IM, with an overshoot of ΔΨm to a level higher than that of the steady state, as was seen in cells upon the depletion of F_o_ subunit ATPTB1 or treatment with the F_o_ proton pore inhibitor oligomycin (30, 36, 53). We therefore measured the ΔΨm of uninduced and induced subunit *b* RNAi cells with the ΔΨm-sensitive dye tetramethylrhodamine ethyl ester (TMRE) by flow cytometry (Fig. 5B). By day 1 post induction, there was a significant increase in the ΔΨm, a level which remained high at day 2 post induction, and then decreased by day 3 to a level 75% of that of uninduced cells.

The hyperpolarisation of the mitochondrial IM can cause the production of reactive oxygen species (ROS) (54–56). We therefore measured the production of mitochondrial superoxide and cellular ROS in subunit *b* RNAi cells before and after induction using flow cytometry and the ROS-sensitive dyes MitoSOX and CM-H_2_DCFHDA respectively. We detected a significant transient increase in mitochondrial superoxide production by day 1 post induction (Fig. 5C), and an increase in cellular ROS production by day 2 post induction (Fig. 5D). Mitochondrial superoxide levels immediately peaked and then decreased over time after subunit *b* depletion, presumably due to the rewiring of electrons from the respiratory complexes contributing to ΔΨm towards the non-proton pumping AOX (30, 57–59). However, cellular ROS molecules continued to accumulate, in a similar fashion to that seen upon the depletion of F_o_ subunit ATPTB1 (30), a culmination of the non-reversible damage caused by subunit *b* depletion, and likely a direct effector of the growth phenotype.

Finally, we investigated whether the growth retardation caused by the lack of subunit *b* is accompanied by disturbance of the cell cycle. Fig. 5E and Fig. 5F show that this in indeed was the case. Analysis of the DNA content of these cells by fluorescence microscopy of 4’, 6-diamidino-2-phenylindole (DAPI) stained cells (Fig. 5E and Fig. 5F, left graph) and flow cytometry analysis of propidium iodide (PI) stained cells (Fig. 5F, right graph) showed a defect in cell cycle progression concomitant with the growth phenotype. After day 2 post induction, a reduction in the proportion of cells going through kinetoplast division (2K1N cells, S stage of the cell cycle, square symbols in Fig. 5F) was seen, with a large decrease in the proportion of cells beginning the cell cycle seen between days 4 and 6 (1K1N, G1, circle symbols in Fig. 5F). There was also an increase in the proportion of zoid cells with no nucleus (marked with I in Fig. 5E, diamond symbols in Fig 5F, incorporated into the “others” category in Fig. 5F left panel), and cells with large or multiple nuclei (marked with II in Fig. 5E, diamond symbols in Fig. 5F, >G2/M and incorporated in the “others” category). Moreover, across this time course approximately 50% of all cells had an aberrant DNA content by day 6. Thus, subunit *b* is required for normal growth and cell cycle progression in procyclic *T. brucei*. However, there are no massive changes in mitochondrial morphology after depletion of subunit *b* in induced cells (Fig. 5E, bottom panel).

## DISCUSSION

The *T. brucei* F_1_F_o_ ATP synthase peripheral stalk subunit *b* could not be identified by sequence homology. This is not that surprising since closely related organism *E. gracilis*, which has an F_1_F_o_ ATP synthase with similarity in architecture and subunit composition to the *T. brucei* version (13, 23, 33, 34), has a highly diverged subunit *b* (14). The peripheral stalk of the *E. gracilis* complex has an extended OSCP which unusually interacts with the central stalk subunit γ, and with a divergent extended subunit *d* which unlike its opisthokont counterpart has a TMD (21, 60). The euglenozoan-specific ATPTB3 and ATPTB4 stabilise the interaction between subunit *d* and OSCP and facilitate an interaction with the catalytic core of the F_1_ moiety via one copy of p18 (13). *E. gracilis* subunit *b* could not be detected by sequence homology but was identified based on its structural similarity, position and topology within the structure. It contains one TMD and is truncated in comparison to its yeast counterpart. Moreover, unlike subunit *b* in other complexes, it does not interact directly with OSCP, the dimerisation interface or subunit 8. Instead, it interacts extensively with subunit *d* along the external peripheral stalk, and with subunits *a* and *f* in the membrane-embedded region.

Interestingly, the *T. brucei* F_1_F_o_ ATP synthase has an extended OSCP as well as a highly diverged extended subunit *d* similar to that of *E. gracilis*. Based on the structure of the peripheral stalk of *E. gracilis* and because no subunit *b* could be identified in *T. brucei*, the question of whether the trypanosomal version of the protein could be either further reduced in length or even be completely absent was raised (14).

We can now answer this question. We have identified a 17 kDa mitochondrial IM protein containing a single predicted TMD close to the N-terminus which is essential for normal growth of procyclic *T. brucei*. The tagged protein pulls down essentially all known F_1_F_o_ ATP synthase subunits and co-migrates with the monomer of the F_1_F_o_ ATP synthase when analysed by BN-PAGE. Ablation of the protein preferentially destabilises F_o_ but not F_1_ ATP synthase subunits and results in a downregulation of the F_o_ moiety-containing complexes on BN gels. RNAi analysis shows that the protein is involved in oxidative phosphorylation and ΔΨm maintenance in a similar way to that previously described for other trypanosomal F_o_ ATP synthase subunits. Finally, HHpred analysis reveals structural similarities to F_1_F_o_ ATP synthase subunits *b* of other species. These results indicate that the 17 kDa protein is the so far elusive subunit *b* of the peripheral stalk of the F_1_F_o_ ATP synthase of trypanosomes. Thus, *T. brucei* indeed does contain a highly diverged subunit *b* orthologue that, while truncated, is 30% larger than the *E. gracilis* protein (145 amino acids compared with 112). This suggests that the *T. brucei* ATP synthase peripheral stalk may have a similar global architecture than the one in *E. gracilis*, although the longer trypanosomal subunit *b* could potentially show unique subunit interactions.

Interestingly, the *T. brucei* subunit *b* has not been detected in previous pull-down MS analyses of the *T. brucei* F_1_F_o_ complexes (33, 61). This could be explained by complex disruption during extraction and/or by the low molecular mass and the few tryptic cleavage sites of subunit *b* which may make detection by proteomic methods difficult (62). However, *T. brucei* subunit *b* had previously been detected in a pull-down analysis of the mitochondrial calcium uniporter, along with 19 other subunits of the ATP synthase (63).

Knockdown of subunit *b* led to a more than 1.5-fold downregulation of nine F_o_ and peripheral stalk subunits, showing that despite its small size and reduced subunit interactions, it is an essential subunit for F_o_ and peripheral stalk assembly and/or for maintenance of their structural integrity. In contrast, all F_1_ subunits remain stable even in the absence of subunit *b*, similar to what has been shown in yeast (64–66), with the F_1_ moiety thereby assembling independently of an intact F_o_ moiety.

Besides many previously identified F_1_F_o_ ATP synthase subunits, our SILAC pulldown and SILAC RNAi analyses recovered a number of trypanosome-specific proteins of unknown function. These proteins, while not the subject of the present study, may well include further as yet unknown trypanosome-specific F_1_F_o_ ATP synthase subunits.

Ablation of trypanosomal ATP synthase subunit *b* results in physiological changes that already have been described for other F_o_ ATP synthase subunits. Interestingly, we also observed a cell cycle phenotype. kDNA replication seems to be inhibited, and possibly as a consequence, the whole cell cycle is disturbed. We did not find any published results reporting the same type of analysis for any other F_1_ or F_o_ ATP synthase subunits of trypanosomes. It is therefore not clear whether ablation of any subunit of the F_1_F_o_ ATP synthase would cause the same phenotype. Indeed, a recent systematic analysis of RNAi cell lines targeting 101 randomly chosen mitochondrial proteins of unknown function in procyclic *T. brucei* showed that ablation of approximately a third of these proteins caused a growth retardation concomitant with abnormalities in cell cycle progression (67). This suggests that an abnormal cell cycle is a common phenotype observed after ablation of mitochondrial proteins that are required for normal growth. However, in the subunit *b* RNAi cell line, the fraction of cells with abnormal N/K configurations was approximately 40% whereas Mbang-Bennet et al. report such strong phenotypes for only three of 37 cell lines showing cell cycle abnormalities. Presently, we do not know the underlying mechanism that causes the cell cycle phenotype detailed here.

In the present study, we have discovered and characterised the perennially elusive subunit *b* of the F_o_ moiety of the trypanosomal ATP synthase. These results, in the absence of a high resolution structure, adds to our understanding of the divergent peripheral stalk in the *T. brucei* enzyme, and will inform the interpretation of future structural studies of the complete trypanosomal ATP synthase complex.

## EXPERIMENTAL PROCEDURES

### Transgenic cell lines

Transgenic *T. brucei* cell lines were generated using the procyclic strain 29–13 (68). Procyclic forms were cultivated at 27°C in SDM-79 (69) supplemented with 10% (v/v) fetal calf serum (FCS) containing G418 (15 μg/ml), hygromycin (25 μg/ml), puromycin (2 μg/ml), and blasticidin (10 μg/ml) as required. RNAi or protein expression was induced in cell lines by adding 1µg/ml tetracycline to the medium.

The Tb927.8.3070 RNAi cell lines were prepared using a pLew100-derived vector containing a blasticidin resistance gene, with the generation of a stem-loop construct occurring by the insertion of the RNAi inserts in opposing directions. The loop is formed by a 460 bp spacer fragment. RNAi plasmids were prepared targeting Tb927.8.3070 via its entire ORF or its entire 3’ UTR (as designated on TriTrypDB). PCR was used to amplify the RNAi targets with primers (F) ACATTAAAGCTTGGATCC<u>ATGGCCTATGTTTCTCCAGC</u> and (R) CGTATTTCTAGACTCGAG<u>CTACTTCGGTAACCGCTGCT</u> (ORF) and (F) ACATTAAAGCTTGGATCC<u>CGGCGGTGCGTGGTTG</u> and (R) CGTATTTCTAGACT CGAG<u>AACGAGAGGAGAGAGACCGC</u> (3’ UTR). RNAi efficiency was verified by RNA extraction and Northern blot, as detailed in (70).

To produce the plasmid for ectopic expression of C-terminal triple c-myc-tagged Tb927.8.3070, the complete ORF was amplified by PCR using primers (F) ACATTAAAGCTT<u>ATGGCCTATGTTTCTCCAGC</u> and (R) CGTATTGGATCC<u>CTTCGGTAACCGCTGCTGAT</u>. The PCR product was cloned into a modified pLew100 vector which contains a puromycin resistance gene as well as a triple epitope tag. Protein expression was verified by SDS-PAGE and immunoblotting of cell lysates.

### Subcellular localisation

The subcellular localisation of Tb927.8.3070 was analysed by generating crude mitochondrial-enriched fractions. 10^8^ cells were incubated in 0.6 M sorbitol, 20 mM Tris-HCl pH 7.5, 2 mM EDTA pH 8 containing 0.015% (w/v) digitonin on ice for 10 min to solubilise the cell membranes. Centrifugation for 5 min at 6,800 g at 4°C yielded a supernatant that is enriched for cytosolic proteins and a crude mitochondrial extract pellet. 2 × 10^6^ cell equivalents of each fraction were analysed by SDS-PAGE and Western.

The mitochondria-enriched pellet was resuspended in 100 mM Na_2_CO_3_ pH 11.5, incubated on ice for 10 min and centrifuged for 10 min at 100’000 g, 4°C to differentiate soluble or loosely membrane associated proteins from integral membrane proteins. 2 × 10^6^ cell equivalents of each fraction were analysed by SDS-PAGE and Western blotting.

Commercially available antibodies were used as follows: mouse c-Myc (Invitrogen; 1:2,000) and mouse EF1a (Merck Millipore; 1:10,000). The polyclonal VDAC (1:1,000) (71) and cytochrome *c* (1:100) (42) antibodies previously produced in our lab were also used. Secondary antibodies for immunoblot analysis were IRDye 680LT goat anti-mouse, and IRDye 800CW goat anti-rabbit (both LI-COR Biosciences, 1:20,000).

For immunofluorescence analysis, cells were fixed with 4% paraformaldehyde in PBS, permeabilised with 0.2% Triton-X100 in PBS and blocked with 2% BSA. Primary antibodies used were mouse anti-c-Myc (1:50) and rabbit anti-ATOM40 (1:1,000), and secondary antibodies were goat anti-mouse Alexa Fluor 596 and goat anti-rabbit Alexa Fluor 488 (both ThermoFisher Scientific; 1:1000).

Slides were mounted with VectaShield containing DAPI (Vector Laboratories). Images were acquired with a DFC360 FX monochrome camera (Leica Microsystems) mounted on a DMI6000B microscope (Leica Microsystems). Images were analysed using LAS AF software (Leica Microsystems) and ImageJ.

### BN Page and quantification

Mitochondrial-enriched pellets from 10^8^ cells/sample were incubated for 15 min on ice in 20 mM Tris-HCl pH 7.4, 50 mM NaCl, 10% glycerol, 0.1 mM EDTA, 1 mM PMSF containing 1% (w/v) digitonin to solubilise mitochondrial membranes. After centrifugation for 15 min at 20’817 g, 4°C, the supernatants were separated on 4–13% gradient gels. The gel was then incubated in SDS-PAGE running buffer (25 mM Tris, 1 mM EDTA, 190 mM glycine, 0.05% (w/v) SDS) to aid the transfer of proteins to the membrane. Quantification of bands was performed using ImageJ.

### ATP production assay

ATP production was measured using substrates succinate, pyruvate, and α-ketoglutarate as described (52). The AAC inhibitor atractyloside was added to samples as a negative control. ATP Bioluminescence assay kit CLS II (Roche Applied Science) was used to measure the ATP concentration of the samples in a luminometer plate reader.

### Flow cytometry

The TMRE Mitochondrial Membrane Potential kit (Abcam) was used to measure ΔΨm. 1 × 10^6^ cells from each sample were resuspended in 1 ml media. All samples were pre-incubated with or without 20 μM uncoupler carbonyl cyanide-*4*-(trifluoromethoxy)phenylhydrazone (FCCP) to provide a negative control for 10 min at 27°C, supplemented with 100 nM TMRE, and left at 27°C for 20 min. Cells were pelleted at 2000 g for 5 min in 5 ml polystyrene round bottom tubes (BD Falcon 352052), and washed three times in 5 ml 0.2% BSA in PBS-G containing 6 mM glucose. Cell pellets were resuspended in 500 μl 0.2% BSA in PBS-G, left for 30 min in foil, and analysed was performed on Novocyte instrument (Agilent), using the 488 nm laser for excitation and detection using the B586/20nm filter. Data was analysed using FlowJo.

For the measurement of ROS production, 3 × 10^6^ cells/sample were analysed. The same protocol as above was used, but cells were supplemented with either 5 μM MitoSOX (Thermo Fisher Scientific) or 10 μM CM-H_2_DCFDA (Thermo Fisher Scientific). The 488 nm laser was used for excitation, with detection using the B586/20nm filter for MitoSOX and the B530/30nm filter for CM-H2DCFDA.

For cell cycle analysis, 1 × 10^6^ cells/sample were fixed with 70% ice cold methanol for 10 minutes, washed in PBS twice and stained with 10 μg/ml PI plus 10 μg/ml RNase A. After incubation for 45 minutes, the samples were analysed using the 488 nm laser for excitation and detection using the B586/20nm filter.

### SILAC-based proteomics and IP experiments

Cells were washed in PBS and taken up in SDM-80 supplemented with 5.55 mM glucose, either light (^12^C_6_/^14^N_χ_) or heavy (^13^C_6_/^15^N_χ_) isotopes of arginine (1.1 mM) and lysine (0.4 mM) (Euroisotop) and 10% dialysed FCS (BioConcept, Switzerland). To guarantee complete labelling of all proteins with heavy amino acids, the cells were cultured in SILAC medium for 6–10 doubling times.

The Tb927.8.3070 RNAi cell line was induced with tetracycline for 3d. 1 × 10^8^ uninduced and 1 ×10^8^ induced cells were harvested and mixed. Crude mitochondria-enriched pellets were obtained by incubating 2 × 10^8^ cells on ice for 10 min in 0.6 M sorbitol, 20 mM Tris-HCl pH 7.5, 2 mM EDTA pH 8 containing 0.015% (w/v) digitonin, and centrifugation (5 min/6,800 g/4°C). The digitonin-extracted mitochondria-enriched pellets generated from these mixed cells were then analysed.

For SILAC-IP experiments, cells were induced for 1d. 1 × 10^8^ uninduced and 1 × 10^8^ induced cells were harvested in triplicate, mixed and subjected to a CoIP protocol as follows.

2 × 10^8^ cells were solubilised for 15 min on ice in 20 mM Tris-HCl pH7.4, 0.1 mM EDTA, 100 mM NaCl, 25 mM KCl containing 1% (w/v) digitonin and 1X Protease Inhibitor mix (Roche, EDTA-free). After centrifugation (15 min, 20000 g, 4°C), the lysate (IN) was transferred to 50 μl of bead slurry, which had been previously equilibrated with respective lysis buffer. The bead slurries used were c-myc-conjugated (EZview red rabbit anti-c-myc affinity gel, Sigma). After incubating at 4°C for 2 hr, the supernatant containing the unbound proteins was removed, the bead slurry was washed three times with lysis buffer and the bound proteins were eluted by boiling the resin for 10 min in 2% SDS in 60 mM Tris-HCl pH 6.8 (IP). Pull down of the bait was confirmed by SDS-PAGE and Western blotting, with 5% of both the input and the flow through samples and 50% of the IP sample loaded.

All SILAC experiments were performed in three biological replicates including a label-switch and analysed by liquid chromatography-mass spectrometry (LC-MS).

### LC-MS and data analysis

Samples generated in Tb927.8.3070 SILAC RNAi experiments were processed for LC-MS analysis (including reduction and alkylation of cysteine residues, tryptic in-solution digestion) as described before (41). Eluates of Tb927.8.3070 SILAC-IP experiments were loaded onto an SDS gel and electrophoresis was performed until the proteins had migrated into the gel for approximately 1 cm. Proteins were visualised using colloidal Coomassie Blue, protein-containing parts of the gel were excised en bloc and cut into smaller cubes, followed by reduction and alkylation of cysteine residues and tryptic in-gel digestion as described before (41).

LC-MS analyses of tryptic peptide mixtures were performed using an Orbitrap Elite mass spectrometer (Thermo Fisher Scientific, Bremen, Germany) connected to an UltiMate 3000 RSLCnano HPLC system (Thermo Fisher Scientific, Dreieich, Germany). Peptides were loaded and concentrated on nanoEase(tm) M/Z Symmetry C18 precolumns (20 mm x 0.18 mm; flow rate, 10 µl/min; Waters) and separated using a nanoEase(tm) M/Z HSS C18 T3 analytical column (250 mm x 75 µm; particle size, 1.8 µm; packing density, 100 Å; flowrate, 300 nl/min; Waters). A binary solvent system consisting of 0.1% formic acid (solvent A) and 30% acetonitrile/50% methanol/0.1% formic acid (solvent B) was used. Peptides of all experiments were loaded and concentrated for 5 min at 7% B, followed by peptide elution applying the following gradients: 7 - 60% B in 295 min, 60 - 95% B in 35 min and 5 min at 95% B (SILAC RNAi) or 7 - 50% B in 105 min, 50 - 95% B in 45 min and 5 min at 95% B (SILAC-IP experiments).

Mass spectrometric data were acquired in data-dependent mode applying the following parameters: mass range of *m/z* 370 to 1,700, resolution of 120,000 at *m/z* 400, target value of 1 × 10^6^ ions, and maximum injection time of 200 ms for MS survey scans. A TOP25 method was used for low energy collision-induced dissociation of multiply charged peptides in the linear ion trap at a normalized collision energy of 35%, an activation q of 0.25, an activation time of 10 ms, a target value of 5,000 ions, a maximum injection time of 150 ms, and a dynamic exclusion time of 45 s.

MaxQuant/Andromeda (version 1.6.5.0, (72, 73)) was used for protein identification and SILAC-based relative quantification. Database searches were performed using the proteome of *T. brucei* TREU927 downloaded from the TriTryp database (https://tritrypdb.org; version 8.1, containing 11,067 entries) and MaxQuant default settings with the exception that one unique peptide was sufficient for protein identification. Carbamidomethylation of cysteine residues was set as fixed modification, N-terminal acetylation and oxidation of methionine were considered as variable modifications, and Arg10 and Lys8 were set as heavy labels. The options ‘requantify’ and ‘match between runs’ were enabled. SILAC ratios were calculated based on unique peptides and at least one ratio count. Results of protein identification and quantification are provided in the Supporting Information as Tables S1 (Tb927.8.3070 SILAC-IP data) and S2 (Tb927.8.3070 SILAC RNAi data).

## ACKNOWLEDGMENTS

We thank Bettina Knapp for assistance in LC-MS analyses and Elke Horn for technical assistance. In the lab of B. W., the study was supported by the Deutsche Forschungsgemeinschaft (DFG, German Research Foundation) project ID 403222702/SFB 1381. In the lab of A. S., the study was supported by the NCCR “RNA & Disease” and in part by grants 175563, both funded by the Swiss National Science Foundation.

## SUPPORTING INFORMATION

**Fig. S1.**
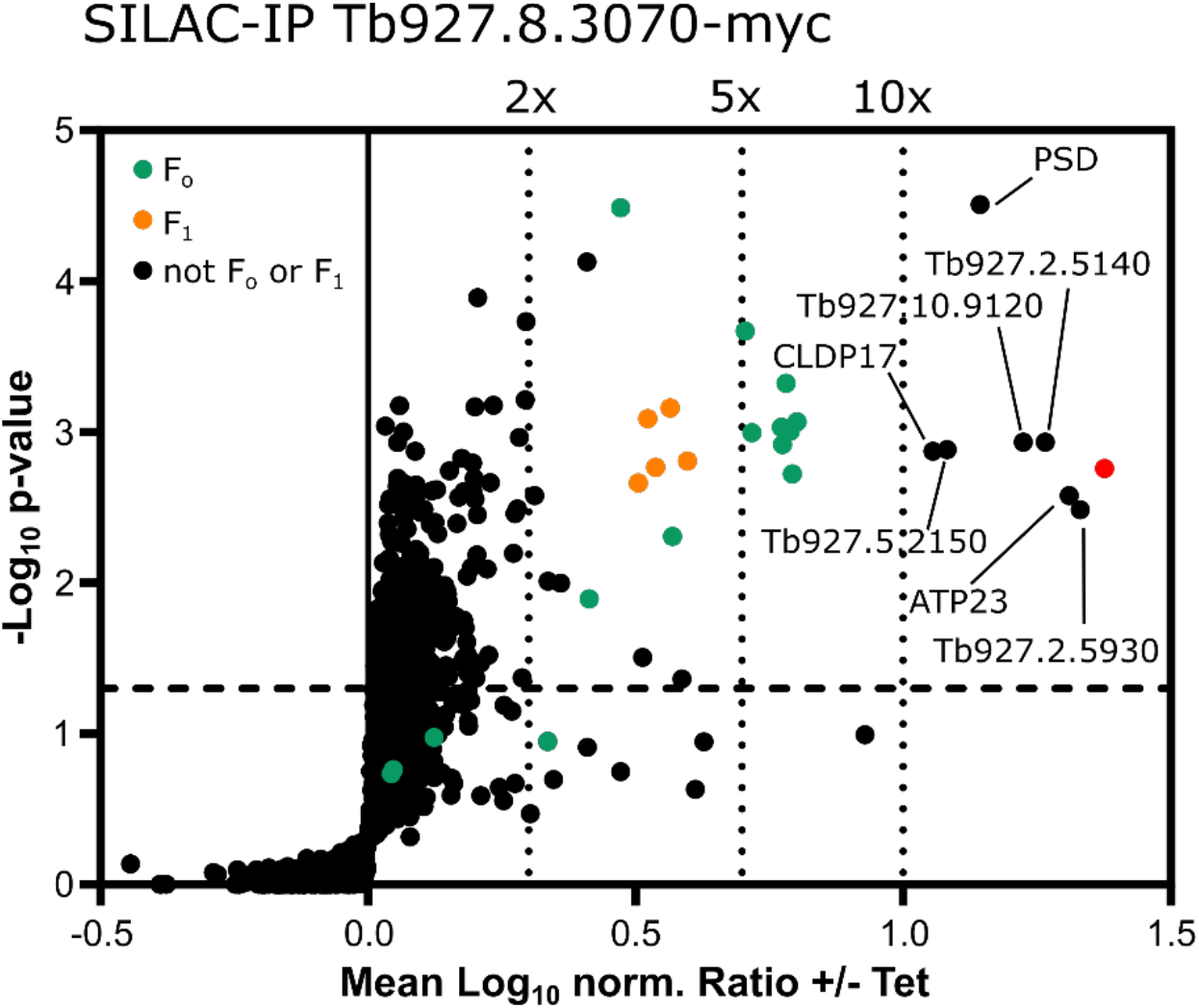
Further proteins that interact with Tb927.8.3070. A volcano plot depicting of the SILAC-IP analysis of crude mitochondrial extracts from Tb927.8.3070-myc expressing cells as shown in Fig 1C. Proteins more than 10-fold enriched are labelled with either their name or accession numbers.

**Fig. S2.**
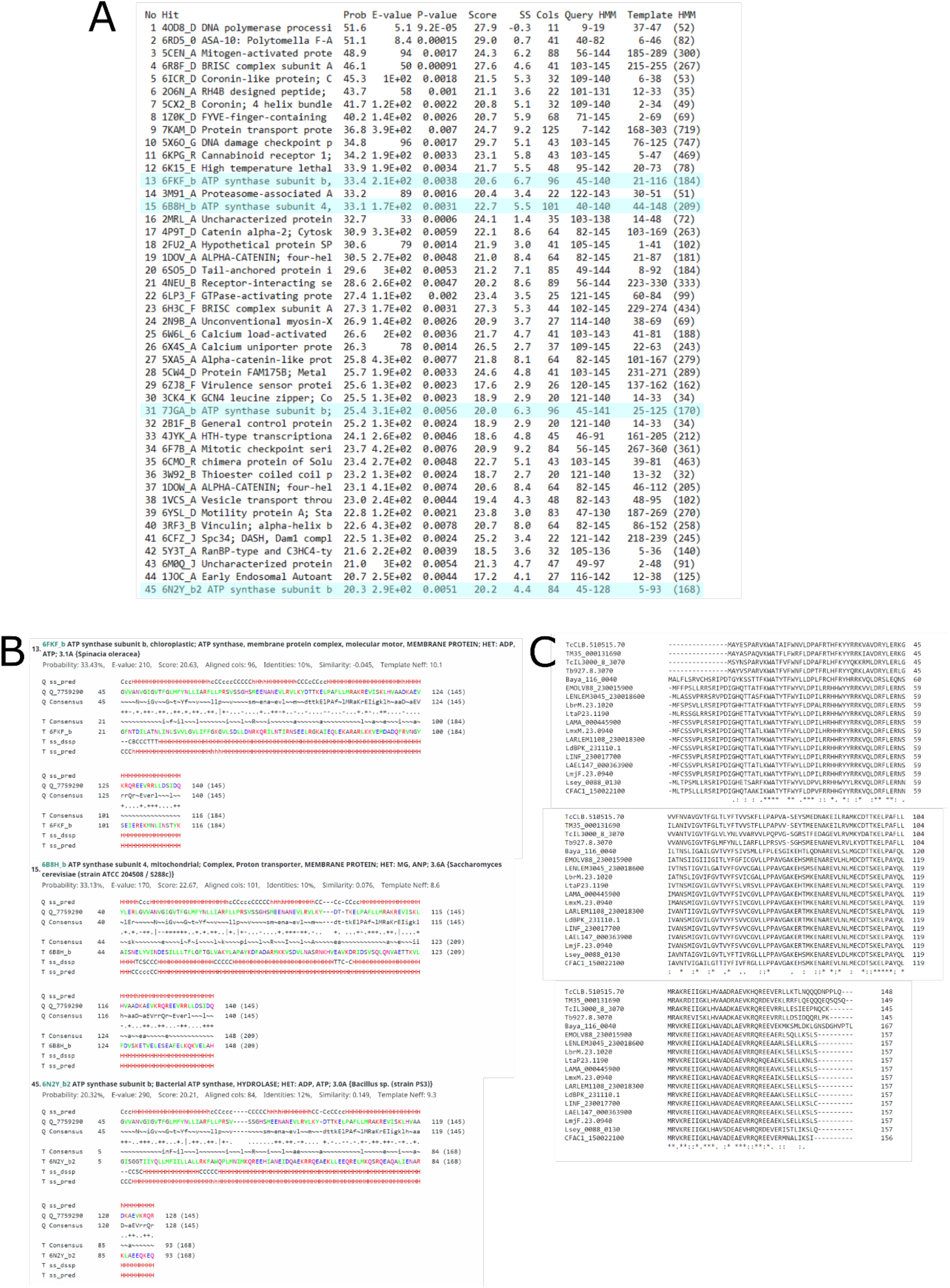
*In silico* analysis of Tb927.8.3070. **(A)** List of HHpred results using Tb927.8.3070 as the input sequence. The hits related to ATP synthase subunit *b* are highlighted in blue. **(B)** The sequence of Tb927.8.3070 that displays secondary structure homology to regions in the ATP synthase subunit *b* of spinach (*Spinacia oleracea*) chloroplasts, yeast (*S. cerevisiae*) and *Bacillus* species using HHpred. **(C)** Sequence alignment between Tb927.8.3070 and its orthologs in Kinetoplastid species using Clustal Omega (78). TcCLB *T. cruzi*, TM *T. theileri*, TcIL *T. congolense*, Baya *B. ayalai*, EMOLV *E. monterogeii*, LENLEM *L. enriettii*, Lbr *L. braziliensis*, Lta *L. tarentolae*, LAMA *L. amazonensis*, Lmx *L. mexicana*, LARLEM *L. arabica*, Ld *L. donovani*, LINF *L. infantum*, LAEL *L. aethiopica*, Lmj *L. major*, Lsey *L. seymouri*, CFAC *C. fasciculata*.

